# Single-value scores of memory-related brain activity reflect dissociable neuropsychological and anatomical signatures of neurocognitive aging

**DOI:** 10.1101/2022.02.04.479169

**Authors:** Anni Richter, Joram Soch, Jasmin M. Kizilirmak, Larissa Fischer, Hartmut Schütze, Anne Assmann, Gusalija Behnisch, Hannah Feldhoff, Lea Knopf, Matthias Raschick, Annika Schult, Constanze I. Seidenbecher, Renat Yakupov, Emrah Düzel, Björn H. Schott

## Abstract

Memory-related functional magnetic resonance imaging (fMRI) activations show age-related differences across multiple brain regions that can be captured in summary statistics like single-value scores. Recently, we described two single-value scores reflecting deviations from prototypical whole-brain fMRI activity of young adults during novelty processing and successful encoding. Here, we investigate the brain-behavior associations of these scores with age-related neurocognitive changes in 153 healthy older adults. All scores were associated with episodic recall performance. The memory network scores, but not the novelty network scores, additionally correlated with medial temporal gray matter and a composite measure comprising pro-active inhibition, episodic memory, tonic alertness, flexibility, and working memory. Our results thus suggest that novelty-network-based fMRI scores show high brain-behavior associations with episodic memory and that encoding-network-based fMRI scores additionally capture individual differences in global cognitive function. More generally, our results suggest that single-value scores of memory-related fMRI provide a comprehensive measure of individual differences in network dysfunction that may contribute to age-related cognitive decline.

## 1. Introduction

Even healthy older adults commonly exhibit a certain degree of cognitive decline and brain structural alterations (Anthony & Lin, 2018; Cabeza et al., 2018; X. Li et al., 2020). While age-related decline of cognitive functions and particularly explicit memory is common, some individuals age more “successfully”, showing comparably preserved memory capability even in advanced age (Nyberg & Pudas, 2019). On the other hand, for example, individuals at risk for Alzheimer’s disease (AD) exhibit accelerated cognitive aging well before clinical onset of the disease. Valid and comprehensive markers of cognitive and functional impairment could facilitate the assessment of age-related neurocognitive changes and provide valuable information about an individual’s extent of brain aging (Frisoni et al., 2017; Jack et al., 2013; Partridge et al., 2018; Tsapanou et al., 2019). As suggested by Hedden et al. (2016), markers that rely on age-related alterations of brain structure and function can be referred to as brain markers or, if obtained using imaging techniques, as imaging biomarkers. Examples include differences in gray matter volume (GMV) (Diaz-de-Grenu et al., 2011; Minkova et al., 2017), white matter (WM) lesion load (Arvanitakis et al., 2016; Tsapanou et al., 2019), memory-related functional magnetic resonance imaging (fMRI) (E. Duzel et al., 2011; Grady & Craik, 2000; Maillet & Rajah, 2014; Soch et al., 2021a), and electrophysiological measures (Babiloni et al., 2020). Other indicators of successful versus accelerated cognitive aging are disease markers, which encompass, among others, positron emission tomography (PET) measures of beta-amyloid (Aβ) and tau deposition (Knopman et al., 2019), but also neuropsychological markers like global cognition, executive function, and episodic memory as assessed with neuropsychological tests (Hassenstab et al., 2015).

Previous studies show that, compared to young individuals, older adults exhibited lower activations of inferior and medial temporal structures and reduced deactivations in the Default Mode Network (DMN) during novelty processing and successful long-term memory encoding (Maillet & Rajah, 2014; Soch et al., 2021a; Billette et al., 2022; E. Duzel et al., 2022). To capture age-related deviations from the prototypical fMRI activations in younger participants, we have previously proposed the use of reductionist fMRI-based scores:

I. The FADE score (*Functional Activity Deviations during Encoding* (E. Duzel et al., 2011), which reflects the *difference* of activations outside and inside a mask representing prototypical activations in a young reference sample, and
II. the SAME score (*Similarity of Activations during Memory Encoding* (Soch et al., 2021a), which reflects the *similarity* of an older adult’s brain response with activation – and also deactivation – patterns in young subjects, adjusted for the between-subjects variance within the young reference sample.

Both markers constitute single-value scores and can be computed either from fMRI novelty (novel vs. highly familiarized images) or subsequent memory contrasts (based on a subsequent recognition memory rating of the to-be-encoded images). They thus constitute reductionist measures of age-related processing differences in either novelty detection or successful encoding, which engage overlapping, but partly separable neural networks (Maass et al., 2014; Soch et al., 2021a; Soch et al., 2021b), with novelty detection not directly translating to encoding success (Poppenk et al., 2010). Scores based on novelty detection versus encoding success may thus indicate age-related deviations in at least partly different cognitive domains. The FADE and SAME scores have previously been associated with memory performance in the encoding task they were computed from (E. Duzel et al., 2011; Soch et al., 2021a), but it is yet unclear whether this relationship is also found with independent, classical neuropsychological assessments of memory. Furthermore, it is not yet known whether the scores are specifically related to hippocampus-dependent memory performance or rather global cognitive function in old age.

Here, we investigate brain-behavior associations of the scores with age-related differences in episodic memory and hippocampal function, as reflected by correlations with memory performance measures and medial temporal lobe (MTL) gray matter volume (GMV), as well as their relationship with other cognitive domains and age-related differences in brain morphology beyond the MTL. To evaluate which neurocognitive functions (hippocampus-dependent memory vs. other cognitive tasks) are significantly related to the four fMRI-based single-value scores (i.e. FADE vs. SAME, obtained from novelty vs. memory contrast) and specifically to age-related differences, we assessed their associations with multiple measures of cognitive ability and of structural brain integrity in a large cross-sectional cohort of healthy older adults. First, we computed correlations between the imaging scores to assess their potential dependence or orthogonality. We then performed multiple regression analyses to test their relationship with performance in different memory tests and other psychometric tasks covering a wide range of cognitive functions. Finally, we assessed associations between the imaging scores and brain morphometric measures (local GMV, WM lesion volume). For an overview of our approach, see Figure 1.

**Figure 1.**
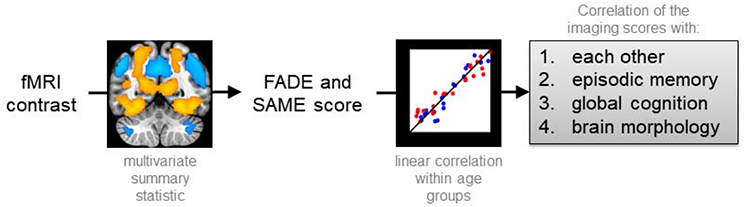
Overview of our approach to investigate the brain-behavior associations of single-value fMRI-based scores with cognitive ability in older adults. Imaging scores were calculated from a voxel-wise fMRI contrast map (warm colors indicate positive effects and cool colors indicate negative effects) and correlated with each other, with neuropsychological test performance in episodic memory, with global cognition, and with measures of brain morphology separately for each age group (red: young, blue: older subjects). All activation maps are superimposed on the MNI template brain provided by MRIcroGL (https://www.nitrc.org/projects/mricrogl/). Figure adapted from Soch et al., 2021a.

## 2. Methods

### 2.1. Participants

The previously described study cohort (Soch et al., 2021a; Soch et al., 2021b) consisted of 259 healthy adults, including 106 young (47 male, 59 female, age range 18-35, mean age 24.12 ± 4.00 years), 42 middle-aged (13 male, 29 female, age range 51-59, mean age 55.48 ± 2.57 years) and 111 older (46 male, 65 female, age range 60-80, mean age 67.28 ± 4.65 years) participants. Additionally, a replication cohort of 117 young subjects (Assmann et al., 2021; 60 male, 57 female, age range 19-33, mean age 24.37 ± 2.60 years) served for outlier detection and a linear discriminant analysis (LDA). Please note that, while this study is based on the same participant sample as previously described, all analyses and results reported in this study have not been published elsewhere. As we found no significant differences between middle-aged and older participants for any of the imaging scores (two-samples *t*-tests: all *p* > .123; also see Soch et al., 2021a, Figure S2), we combined them into one age group to increase sample size and thus the statistical power of the analyses performed here (N = 153, 59 male, 94 female, age range 51-80, mean age 64.04 ± 6.74 years).

According to self-report, all participants were right-handed, had fluent German language skills and did not take any medication for neurological or mental disorders. A standardized neuropsychiatric interview was used to exclude present or past mental disorder, including alcohol or drug dependence.

Participants were recruited via flyers at the local universities (mainly the young subjects), advertisements in local newspapers (mainly the older participants) and during public outreach events of the institute (e.g., *Long Night of the Sciences*).

Data were collected at the Leibniz Institute for Neurobiology in Magdeburg in collaboration with the German Center for Neurodegenerative Diseases in Magdeburg and the Otto von Guericke University of Magdeburg as part of a project within the *Autonomy in Old Age* research alliance. All participants gave written informed consent in accordance with the Declaration of Helsinki (World Medical Association, 2013) and received financial compensation for participation. The study was approved by the Ethics Committee of the Faculty of Medicine at the Otto von Guericke University of Magdeburg.

### 2.2. Neuropsychological assessment

We conducted a number of common psychometric tests that cover a wide range of psychological constructs like attention, different aspects of memory, including short- and long-term memory, working memory as well as executive functions, such as interference control and flexibility. The tests are described in detail in the Supplementary Material; the variables and psychological constructs are summarized in Table 1. Additionally, the Multiple-Choice Vocabulary Test (MWT-B; Lehrl, 2005) was performed as a proxy for crystallized verbal intelligence. It consists of 37 items with increasing difficulty, each item containing one real word and four verbally similar but meaningless pseudo-words of which the participant has to mark the correct one. Data were collected using custom code written in Presentation (0.71, Neurobehavioral Systems, www.neurobs.com).

**Table 1.** Tests and variables of the neuropsychological testing battery.

| test | variables | psychological construct | young subjects<br>M ± SD (N) | older subjects<br>M ± SD (N) | statistics |
| --- | --- | --- | --- | --- | --- |
| Verbal Learning and Memory Test (VLMT) | number of correctly named words of:<br>- repetitions of list A (sum score)<br>**- distractor list B**<br>- recall of list A<br>- 30-min delayed recall of list A<br>**- one-day delayed recall of list A** | learning ability<br>pro-active inhibition<br>retro-active inhibition<br>episodic memory<br>episodic memory | 67.02 ± 6.09 (102)<br>10.14 ± 2.68 (103)<br>14.47 ± 1.02 (103)<br>14.44 ± 1.09 (104)<br>13.94 ± 1.49 (100) | 53.42 ± 9.38 (152)<br>6.36 ± 2.02 (152)<br>11.32 ± 2.84 (152)<br>11.43 ± 2.88 (152)<br>9.26 ± 3.43 (148) | $t = 14.01, p < .001$<br>$t = 12.17, p < .001$<br>$t = 12.53, p < .001$<br>$t = 11.74, p < .001$<br>$t = 14.70, p < .001$ |
| Logical Memory subtest from the WMS | number of story details retrieved at:<br>- immediate recall<br>- 30-min delayed recall<br>- one-day delayed recall | learning ability<br>episodic memory<br>episodic memory | 31.35 ± 7.32 (103)<br>29.85 ± 7.99 (103)<br>29.21 ± 7.77 (102) | 25.45 ± 6.27 (149)<br>22.99 ± 6.58 (148)<br>21.93 ± 6.83 (146) | $t = 6.66, p < .001$<br>$t = 7.18, p < .001$<br>$t = 7.63, p < .001$ |
| Alertness subtest from the TAP | reaction on the appearance of a cross:<br>**- RT in trials with cue tone [ms]**<br>- RT in trials without cue tone [ms] | tonic alertness<br>phasic alertness | 249.91 ± 29.71 (102)<br>276.28 ± 30.40 (102) | 295.54 ± 54.87 (144)<br>329.74 ± 58.40 (144) | $t = -8.39, p < .001$<br>$t = -9.34, p < .001$ |
| Flexibility subtest from the TAP | switching attention between targets:<br>- error rate<br>**- RT [ms]** | flexibility<br>flexibility | 4.42 ± 4.62 (102)<br>1146.73 ± 264.59 (101) | 11.25 ± 13.19 (147)<br>2006.76 ± 575.52 (147) | $t = -5.78, p < .001$<br>$t = -15.84, p < .001$ |
| Flanker task | incongruent vs. congruent trials:<br>- RT difference [ms] | interference processing | 111.26 ± 52.52 (103) | 213.37 ± 133.95 (140) | $t = -8.20, p < .001$ |
| N-Back task | responses on reoccurring letters:<br>- 1-back corrected hit rate<br>- 1-back RT [ms]<br>**- 2-back corrected hit rate**<br>- 2-back RT [ms]<br>- 3-back corrected hit rate<br>- 3-back RT [ms] | working memory<br>working memory<br>working memory<br>working memory<br>working memory<br>working memory | 97.45 ± 4.63 (100)<br>433.17 ± 54.23 (100)<br>65.29 ± 28.39 (104)<br>588.91 ± 100.65 (104)<br>23.67 ± 34.40 (103)<br>630.91 ± 118.27 (103) | 89.65 ± 18.16 (139)<br>490.50 ± 86.10 (139)<br>20.35 ± 37.47 (150)<br>663.39 ± 108.36 (150)<br>-11.77 ± 31.03 (149)<br>708.45 ± 150.52 (149) | $t = 4.85, p < .001$<br>$t = -6.30, p < .001$<br>$t = 10.86, p < .001$<br>$t = -5.55, p < .001$<br>$t = 8.52, p < .001$<br>$t = -4.57, p < .001$ |

Bold type: variables that best discriminate between age groups (see the linear discriminant analysis). RT: reaction time. WMS: Wechsler Memory Scale (Härtig et al., 2000). TAP: Test Battery for Attention (Zimmermann & Fimm, 1993). VLMT: Verbal Learning and Memory Test (Helmstaedter et al., 2001).

### 2.3. Subsequent Memory Paradigm for fMRI

During the fMRI subsequent memory experiment, participants performed an incidental visual memory encoding task with an indoor/outdoor judgment (E. Duzel et al., 2018). Subjects viewed photographs showing indoor and outdoor scenes, which were either novel at the time of presentation (44 indoor and 44 outdoor scenes) or were repetitions of two highly familiar “master” images (22 indoor and 22 outdoor trials), one indoor and one outdoor scene pre-familiarized before the actual experiment (Soch et al., 2021b). Thus, during encoding, every subject was presented with 88 unique (i.e. novel) images and 2 master images that were presented 22 times each. Participants were instructed to categorize images as “indoor” or “outdoor” via button press as the incidental encoding task (i.e., participants were unaware that their memory for the pictures would later be tested). Each picture was presented for 2.5 s, followed by a variable delay between 0.70 s and 2.65 s.

Approximately 70 minutes (70.19 ± 3.60 min) after the start of the fMRI session, subjects performed a computer-based recognition memory test outside the scanner, in which they were presented with the 88 images that were shown once during the fMRI encoding phase (*old*) and 44 images they had not seen before (*new*). Participants rated each image on a five-point Likert scale from 1 (“definitely new”) over 3 (“undecided”) to 5 (“definitely old”; for detailed experimental procedure, see Assmann et al., 2021; Soch et al., 2021b). Data were collected using custom code written in Presentation (0.55, Neurobehavioral Systems, www.neurobs.com).

### 2.4. Magnetic Resonance Imaging

Structural and functional MRI data were acquired on two Siemens 3T MR tomographs (Siemens Verio: 58 young, 83 older; Siemens Skyra: 48 young, 70 older), following the exact same protocol as used in the DELCODE study (E. Duzel et al., 2018; Jessen et al., 2018).

A T1-weighted MPRAGE image (TR = 2.5 s, TE = 4.37 ms, flip-α = 7°; 192 slices, 256 × 256 in-plane resolution, voxel size = 1 × 1 × 1 mm) was acquired for co-registration and improved spatial normalization. Phase and magnitude fieldmap images were acquired to improve correction for artifacts resulting from magnetic field inhomogeneities (*unwarping*). Furthermore, a fluid-attenuated inversion recovery (FLAIR) image was acquired (TR = 5.0 s, TE = 395 ms, inversion time = 1.8 s, voxel size = 1 × 1 × 1 mm) and employed for WM lesion quantification.

For functional MRI (fMRI), 206 T2*-weighted echo-planar images (TR = 2.58 s, TE = 30 ms, flip-α = 80°; 47 slices, 64 × 64 in-plane resolution, voxel size = 3.5 × 3.5 × 3.5 mm) were acquired in interleaved-ascending slice order (1, 3, …, 47, 2, 4, …, 46). The total scanning time during the task-based fMRI session was approximately 9 minutes (Soch et al., 2021b).

#### 2.4.1. Neuroimaging single-value scores (FADE and SAME scores)

Using Statistical Parametric Mapping, Version 12 (SPM12; https://www.fil.ion.ucl.ac.uk/spm/software/spm12/, University College London, UK), we generated single-subject contrast images representing effects of novelty processing (by contrasting novel with familiar images) and subsequent memory effects (by parametrically modulating the BOLD response to novel images as a function of later remembering or forgetting). Specifically, the effect of subsequent memory on fMRI activity during encoding was quantified as the mean-centered and arcsine-transformed subject’s response in a subsequent recognition memory test (ranging from 1 to 5).

As described previously (Soch et al., 2021a) the FADE and SAME scores are based on:

I. computing a reference map showing significant activations (and, for the SAME score, additionally significant deactivations) on each of the two fMRI contrasts (i.e. novelty processing or subsequent memory) within young subjects, and
II. calculating summary statistics quantifying the amount of deviation (FADE score) or similarity (SAME score) for a given older subject with respect to the prototypical (de-)activations seen in young subjects.

More precisely, let *J*_+_ be the set of voxels showing a positive effect in young subjects at an *a priori* defined significance level (here: *p* < 0.05, FWE-corrected, extent threshold k = 10 voxels), and let *t_ij_* be the t-value of the *i*-th older subject in the *j*-th voxel on the same contrast. Then, the FADE score of this subject is given by

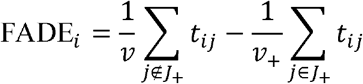

where *v*_+_ and *v* is the number of voxels inside and outside *J*_+_, respectively (Soch et al., 2021a). A larger FADE score signifies higher deviation of an older adult’s memory – or novelty – response from the prototypical response seen in young adults.

Now consider *J*_−_, the set of voxels showing a negative effect in young subjects at a given significance level. Furthermore, let 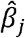 be the average contrast estimate in young subjects, let 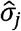 be the standard deviation of young subjects on a contrast at the *j*-th voxel, and let 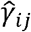 be the contrast estimate of the *i*-th older subject at the *j*-th voxel. Then, the SAME score is given by

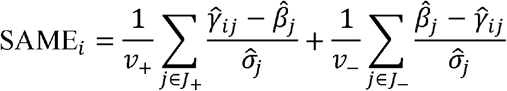

where *v*_+_ and *v*_−_ are the numbers of voxels in *J*_+_ and *J*_−_, respectively (Soch et al., 2021a). Note how the directions of the difference in the two sums are different, in order to accumulate both reduced activations (sum over *J*_+_) and reduced deactivations (sum over *J*_−_). Thus, a higher SAME score indicates higher similarity of an older adult’s brain response with the activation and deactivation patterns seen in young subjects. Simplified, this means that the magnitudes of the SAME (the higher the more similar) and FADE (the higher the less similar) scores have opposing meanings. As further becomes evident from the equation, the SAME score extends the concept underlying the FADE score by:

I. considering deactivation patterns in addition to activation patterns by quantifying reduced deactivations, and
II. accounting for the interindividual variability within the reference sample of young subjects via dividing by their estimated standard deviation.

Hereafter we refer to the scores as follows:

- FADE score computed from the novelty contrast: FADE novelty score
- SAME score computed from the novelty contrast: SAME novelty score
- FADE score computed from the memory contrast: FADE memory score
- SAME score computed from the memory contrast: SAME memory score.

As an initial, exploratory analysis, we computed voxel-wise regressions of the fMRI novelty and subsequent memory contrasts with the imaging scores. Results are reported at *p*_cluster_ < 0.05 using family-wise error rate (FWE) cluster-level correction and an uncorrected cluster-forming threshold of *p*_voxel_< 0.001 (Eklund et al., 2016).

#### 2.4.2. Brain morphometry

VBM analyses were conducted to examine morphological differences of local GMV employing CAT12 using the T1-weighted MPRAGE images. Data processing and analysis were performed as described previously (Assmann et al., 2021; Gvozdanovic et al., 2020; Weise et al., 2019), with minor modifications. Images were segmented into gray matter, WM and cerebrospinal fluid-filled spaces using the segmentation algorithm provided by CAT12. Segmented gray matter images were normalized to the SPM12 DARTEL template, employing a Jacobian modulation and keeping the spatial resolution at an isotropic voxel size of 1 mm^3^. Normalized gray matter maps were smoothed with an isotropic Gaussian kernel of 6 mm at FWHM. Statistical analysis was performed separately for the two age groups using a regression model that included total intracranial volume (TIV) as a covariate. Voxels outside the brain were excluded by employing threshold masking (relative threshold: 0.2) that removed all voxels whose intensity fell below 20% of the mean image intensity (Scarpazza et al., 2015). We computed voxel-wise regressions of the fMRI novelty and subsequent memory contrasts. VBM results are reported at *p*_cluster_ < 0.05 using FWE cluster-level correction and an uncorrected cluster-forming threshold of *p*_voxel_ < 0.001 (Eklund et al., 2016).

Furthermore, we investigated individuals’ brain volumes for WM lesions. Subcortical WM hyperintensities were determined via automatic segmentation in T2-weighted FLAIR images using the Lesion Prediction Algorithm, as implemented in Lesion Segmentation Toolbox (LST v3.0.0; https://www.applied-statistics.de/lst.html) based on the Computational Anatomy Toolbox (CAT12; http://www.neuro.uni-jena.de/cat/, University Hospital Jena, Germany) as described previously (Schmidt et al., 2012; Gaubert et al., 2021). For normalization purposes, WM lesion volume and GMV were divided by the estimated TIV (Guo et al., 2019).

### 2.5. Statistical analysis

Data were analyzed using custom code written in Matlab (2016b) and IBM^®^ SPSS^®^ Statistics, Version 21. We performed step-wise correlational analyses separately for age groups. Firstly, we investigated the potential correlations of the imaging scores among each other. Secondly, we tested their relationship with performance in different memory tests. Thirdly, we correlated the scores with performance in other psychometric tasks covering a wide range of cognitive functions. Finally, we tested for associations between the imaging scores and brain morphometric measures. For an overview of our approach, see Figure 1.

Given the extensive neuropsychological testing battery, which may have included some redundancies (see Table 1), we first aimed to reduce the number of variables to avoid excessive multiple testing. Specifically, we aimed to only include those variables that best separated the age groups. We thus performed a multivariate test of differences using an LDA. A full list of tests and variables included in our LDA can be found in Table 1. To increase the number of young participants, we added the young replication cohort (see Section 2.1) to the analysis, as their neuropsychological assessment included the same cognitive tests. We excluded values that were classified as extreme outliers based on the interquartile range (IQR; × > 3rd quartile + 3*IQR, × < 1st quartile – 3*IQR) in the psychometric tasks separately for each age group (see Supplementary Table S2). We used the step-wise LDA method that stops including tests in the discriminant function (i.e. the linear combination of the performance in the tests that best differentiate between age groups) when there is no longer a significant change in Wilks’ *Λ*. The final set of tests selected with this approach was employed for regression analyses with the SAME and FADE scores. Additionally, we used the composite score gained from the discriminant function as a proxy for global cognition.

For the memory test of the pictures shown during fMRI scanning, memory performance was quantified as A’, the area under the curve (AUC) from the receiver-operating characteristic (ROC) describing the relationship between *false alarms* (“old” responses to new items) and *hits* (“old” responses to previously seen items; see Soch et al., 2021a, Appendix B).

For comparison of age groups, we used paired *t-*tests unless stated otherwise. Whenever Levene’s test was significant, statistics were adjusted, but for better readability, uncorrected degrees of freedom are reported. For the correlational analysis, we used Pearson’s correlations. As the SAME scores can be split into separate components reflecting activations versus deactivations, we performed *post-hoc* correlational analysis with the SAME scores’ activation and deactivation components to unravel possible specific contributions of the components to the significant effects. Whenever appropriate we compared dependent correlation coefficients as described by Meng et al. (1992). We used multiple regression analyses to test the associations of the imaging scores computed from one contrast (novelty vs. memory) as independent variables and the test measures as different dependent variables. As *post-hoc* tests, we used one-sample *t*-tests to examine the unique impact of the coefficients. Significance level was set to *p* < 0.05, two-sided.

## 3. Results

### 3.1. Demographic data

Young and older adults did not differ significantly with respect to gender ratio, ethnic composition or ApoE genotype (*χ*^2^ tests: all *p* > .088, see Table S1). There were significant differences regarding medication, endocrine-related surgeries (e.g. thyroidectomy and oophorectomy), and level of education: 94% of young subjects, but only about 50% of the older subjects had received the German graduation certificate qualifying for academic education (“Abitur”), most likely due to historical differences in educational systems (for a detailed discussion, see Soch et al., 2021a, Supplementary Material). Using the Multiple-Choice Vocabulary Test (MWT-B; Lehrl, 2005), a screening of verbal intelligence, we could confirm that older participants had comparable or even superior verbal knowledge (*z* = − 8.11, *p* < .001), which did not correlate with the imaging scores (all *p* > .203).

Age groups differed significantly for all imaging scores (two-sample *t*-tests: all *p* < .001), except for the FADE score computed from the novelty contrast (*p* = .910; for a discussion, see Soch et al., 2021a).

### 3.2. Voxel-wise representation and inter-correlation of the imaging scores

To help interpreting the subsequently reported results, we computed voxel-wise regressions of the fMRI contrasts with each imaging score for the older adults group. While the FADE score computed from the novelty contrast was rather specifically associated with an occipital and parahippocampal network (Figure S1, upper left part), the FADE score computed from the memory contrast moreover showed positive correlations bilateral with fronto-parietal networks (Figure S1, upper right part). The SAME scores additionally captured a wide range of processes in the DMN (i.e., precuneus and medial prefrontal cortex), which can mainly be attributed to the scores’ negative components. All scores significantly correlated with the contrast they were constructed from (see Figure S1 and Tables S3-6 for details; note that this analysis is partly circular, as the imaging score of each participant were computed from the individual fMRI contrasts. The SAME score computed from the novelty contrast additionally showed a significant positive correlation with the fMRI memory effect in the striatum, precuneus, and middle occipital gyrus (see Figure 2 and Table S7).

**Figure 2.**
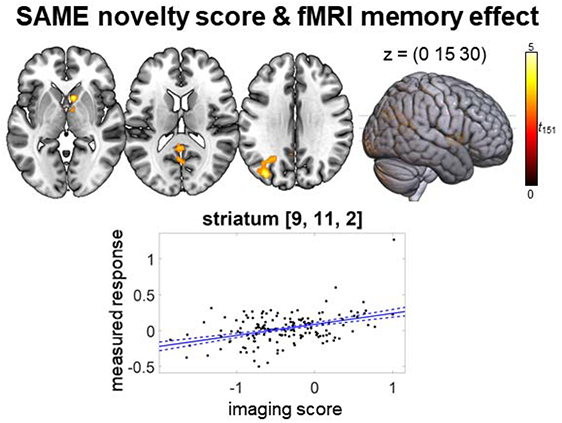
Regression analysis of SAME novelty score and fMRI memory effect (positive effect) in older adults. *p* < .05, family-wise error-corrected at cluster level, cluster-defining threshold *p* < .001, uncorrected. All activation maps are superimposed on the MNI template brain provided by MRIcroGL (https://www.nitrc.org/projects/mricrogl/).

To investigate the scores’ similarity, we correlated them with each other. The scores obtained from the same contrast, that is, novelty or memory, showed significant negative correlations (all *p* < .001; see Figure S2), reflecting the fact that FADE and SAME scores were constructed in opposite ways. Importantly, neither FADE nor SAME scores obtained from the different contrasts (i.e. novelty processing vs. subsequent memory) correlated significantly with each other (*p* > .768), suggesting that they assess different constructs. The remaining correlations were not significant (*p* > .092). *Post-hoc* correlational analysis with the SAME scores’ activation and deactivation components revealed that both components contributed to the correlations with the FADE scores (novelty: activation: *r* = −.646, *p* < .001, deactivation: *r* = −.160, *p* = .048; memory: activation: *r* = −.670, *p* < .001, deactivation: *r* = −.434, *p* < .001). As expected from the construction of the scores, the correlations of the FADE scores with the activation components of the SAME scores were stronger than those with the deactivation components (Fisher’s *z*-test for dependent correlation coefficients: novelty: *z* = − 4.46, *p* <.001, memory: *z* = −2.68, *p* =.007).

### 3.3. The imaging scores are associated with different tests of episodic memory

As the imaging scores were obtained from an fMRI paradigm targeting episodic memory encoding, we first tested for associations with performance in episodic memory tests. These included the recognition memory test of the fMRI experiment itself (70 minutes after onset of the experiment) as well as 30-minutes and one-day delayed recalls of the Verbal Learning and Memory Test (VLMT; Helmstaedter et al., 2001; see supplementary methods) and the Logical Memory subtest from the Wechsler Memory Scale (WMS; Härting et al., 2000). As expected, older participants performed significantly worse in all memory tests compared to young participants (all *p* < .001; see Table 1).

As shown in the previous section, the two sets of imaging scores, FADE novelty and SAME novelty, and FADE memory and SAME memory, are strongly correlated, especially in the case of the memory scores. Thus, while the novelty and memory scores are independent, this is not the case for the SAME and FADE metrics derived from the same contrasts, which share significant fractions of their variances. Therefore, to ascertain the extent to which these metrics explain unique versus shared variance in measures of cognitive performance, we employed a multiple regression approach using the scores derived from the same contrast as independent variables in one model and the memory test measures as different dependent variables.

For the metrics computed from the novelty contrast (see Table 2, upper part, and Figure 3, left side), the scores significantly contributed meaningful information in the explanation

- of memory performance for the pictures shown during fMRI scanning (*F*_2,150_ = 7.62, *p* = .001), with unique impact of the SAME score (FADE: *p* = .098, SAME: *t* =3.79, *p* < .001),
- of performance in the WMS logical memory test 30 minutes delayed recall (*F*_2,145_ = 9.34, *p* < .001), with unique impact of the FADE score (FADE: *t* = −2.84, *p* = .005, SAME: *p* = .434), and
- of performance in the WMS logical memory test one day delayed recall (*F*_2,143_ = 9.21, *p* < .001), with unique impact of the FADE score (FADE: *t* = −2.54, *p* = .012, SAME: *p* = .258).

**Figure 3.**
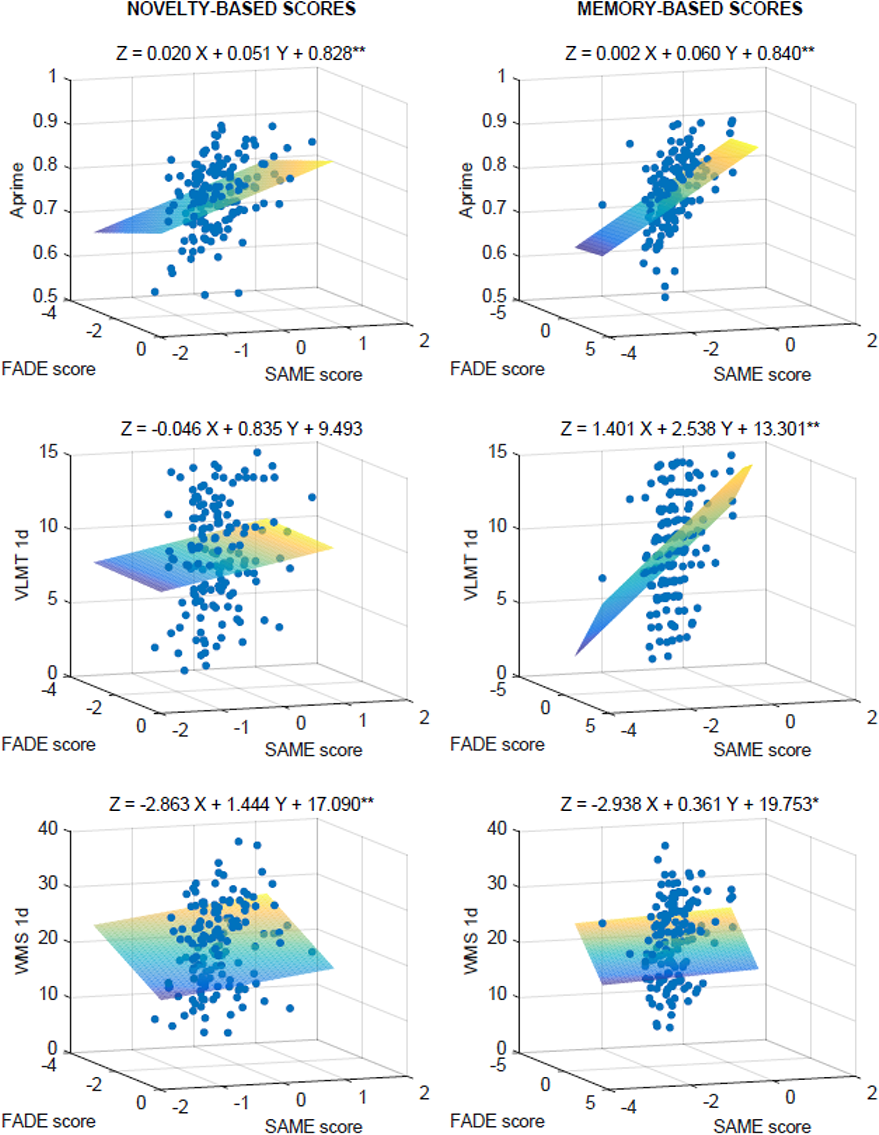
3D-Scatter plots for the regression analyses of FADE vs. SAME scores from the novelty (left) and memory contrast (right) for the memory tests as dependent variables: Aprime, the one-day delayed recall of the VLMT, and the one-day delayed recall of the WMS. * Model is significant at the .05 level. ** Model is significant at the .01 level.

**Table 2.** Regression analyses of the imaging scores with memory tests in older adults.

| **Novelty-based scores** |  |  |  |
| --- | --- | --- | --- |
|  | **Regression model** | **FADE** | **SAME** |
| A' | $R^2 = .092$<br>$F_{2,150} = 7.62^{**}$<br>$p = .001$ | $\beta = 0.020$<br>$t = 1.67$<br>$p = .098$ | $\beta = 0.051$<br>$t = 3.79^{**}$<br>$p < .001$ |
| VLMT<br>30 min | $R^2 = .021$<br>$F_{2,149} = 1.64$<br>$p = .198$ | | |
| VLMT<br>1d | $R^2 = .019$<br>$F_{2,145} = 1.41$<br>$p = .247$ | | |
| WMS<br>30 min | $R^2 = .114$<br>$F_{2,145} = 9.34^{**}$<br>$p < .001$ | $\beta = -3.079$<br>$t = -2.84^{**}$<br>$p = .005$ | $\beta = 0.959$<br>$t = 0.78$<br>$p = .434$ |
| WMS<br>1d | $R^2 = .114$<br>$F_{2,143} = 9.21^{**}$<br>$p < .001$ | $\beta = -2.863$<br>$t = -2.54^*$<br>$p = .012$ | $\beta = 1.444$<br>$t = 1.14$<br>$p = .258$ |
| **Memory-based scores** |  |  |  |
|  | **Regression model** | **FADE** | **SAME** |
| A' | $R^2 = .194$<br>$F_{2,150} = 18.10^{**}$<br>$p < .001$ | $\beta = 0.002$<br>$t = 0.10$<br>$p = .923$ | $\beta = 0.060$<br>$t = 3.30^{**}$<br>$p = .001$ |
| VLMT<br>30 min | $R^2 = .067$<br>$F_{2,149} = 5.35^{**}$<br>$p = .006$ | $\beta = 0.667$<br>$t = 0.79$<br>$p = .434$ | $\beta = 1.833$<br>$t = 2.36^*$<br>$p = .019$ |
| VLMT<br>1d | $R^2 = .068$<br>$F_{2,145} = 5.25^{**}$<br>$p = .006$ | $\beta = 1.401$<br>$t = 1.38$<br>$p = .168$ | $\beta = 2.538$<br>$t = 2.74^{**}$<br>$p = .007$ |
| WMS<br>30 min | $R^2 = .058$<br>$F_{2,145} = 4.45^*$<br>$p = .013$ | $\beta = -2.801$<br>$t = -1.40$<br>$p = .163$ | $\beta = 0.416$<br>$t = 0.23$<br>$p = .818$ |
| WMS<br>1d | $R^2 = .058$<br>$F_{2,143} = 4.37^*$<br>$p = .014$ | $\beta = -2.938$<br>$t = -1.42$<br>$p = .158$ | $\beta = 0.361$<br>$t = 0.19$<br>$p = .849$ |

Note: Number of participants in different tests varies due to completeness of datasets. WMS: Wechsler Memory Scale (Härting et al., 2000). VLMT: Verbal Learning and Memory Test (Helmstaedter et al., 2001). \* Effect is significant at the .05 level. \*\* Effect is significant at the .01 level.

For the metrics computed from the memory contrast (see Table 2, lower part, and Figure 3, right side), the scores significantly contributed meaningful information in the explanation

- of memory performance for the pictures shown during fMRI scanning (*F*_2, 150_ = 18.10, *p* < .001), with unique impact of the SAME score (FADE: *p* = .923, SAME: *t* =3.30, *p* = .001),
- of performance in the VLMT 30 minutes delayed recall (*F*_2, 149_ = 5.35, *p* = .006), with unique impact of the SAME score (FADE: *p* = .434, SAME: *t* = 2.36, *p* = .019),
- of performance in the VLMT one day delayed recall (*F*_2, 145_ = 5.25, *p* = .006), with unique impact of the SAME score (FADE: *p* = .168, SAME: *t* = 2.74, *p* = .007),
- of performance in the 30 minutes delayed recall of WMS logical memory test (*F*_2, 145_ = 4.45, *p* = .013) with no unique impact of either one score (FADE: *p* = .163, SAME: *p* = .818), indicating that their shared variance contributes to the significant association, and
- of performance in the one day delayed recall of WMS logical memory test (*F*_2, 143_ = 4.37, *p* = .014), also with no unique impact of either one score (FADE: *p* = .158, SAME: *p* = .849).

As expected from the construction of the scores, associations with the FADE score (which focuses on deviations from young adults’ prototypical activation patterns) were negative, while associations with the SAME scores (which focus on similarities) were positive. In the group of young adults, no significant regression results were observed (novelty scores: all *p* > .167; memory scores: all *p* > .055).

Next, we explored whether the observed unique associations with the SAME scores were driven by additionally considering deactivations using *post-hoc* multiple regression analyses with the activation and deactivation components as independent variables and one-sample *t*-tests for the coefficients. Indeed, the associations of the SAME novelty score with A’ (activation: *p* = .794, deactivation: *t* = .267, *p* = .001) and of the SAME memory score with VLMT delayed recalls (activation: all *p* > .246, deactivation: all *p* = .006) were carried by the deactivation component. This may be a reason why the FADE novelty score did not correlate with A’, as it did not consider deactivation differences between young and older subjects. The association of the SAME memory score with A’ was driven by both components (activation: *t* = .235, *p* = .004, deactivation: *t* = .329, *p* < .001).

### 3.4. Relationship of the imaging scores with measures of global cognition

To evaluate brain-behavior-associations with the imaging scores beyond hippocampus-dependent memory, we performed regression analyses with neuropsychological tests of other cognitive constructs. Compared to younger participants, older participants showed significantly lower performance in all neuropsychological tests (all *p* < .001; see Table 1). We first computed an LDA to reduce the number of tests and to obtain a proxy for global cognition by including the composite score gained from the discriminant function. Of our 376 subjects (including a young replication sample to increase sample size (see Section 2.1), 107 could not be included in the LDA due to at least one missing value. The final LDA thus included 269 subjects (158 young and 111 older participants). Five variables contributed significantly to the discrimination between age groups as part of the discriminant function (Wilks’ Λ = .348, *p* < .001):

I. the number of words recalled in the distractor trial of the VLMT (standardized canonical discriminant coefficient: .277),
II. the number of words recalled in the one-day delayed recall of the VLMT (.364),
III. the corrected hit rate in the 2-back task (.260),
IV. the reaction time (RT) in the flexibility task (−.478), and
V. the RT of alertness trials with tone (−.225).

90.1 % of the participants could successfully be classified as either young or older when using this discriminant function (young subjects: 92.8 %; older subjects: 86.4 %). We focused our regression analyses on the aforementioned variables best discriminating between age groups, with the exception of the VLMT one-day delayed recall, which was already considered in our analysis of episodic memory tests.

Regarding the metrics computed from the novelty contrast (see Table 3, upper part, and Figure 4, left side), the scores contributed meaningful information in the explanation of performance in the recall of the VLMT distractor list (*F*_2, 149_ = 3.45, *p* = .034), with no unique impact of either one score (FADE: *p* = .100, SAME: *p* = .587), indicating that shared variance of SAME and FADE scores contributes to the significant association.

**Figure 4.**
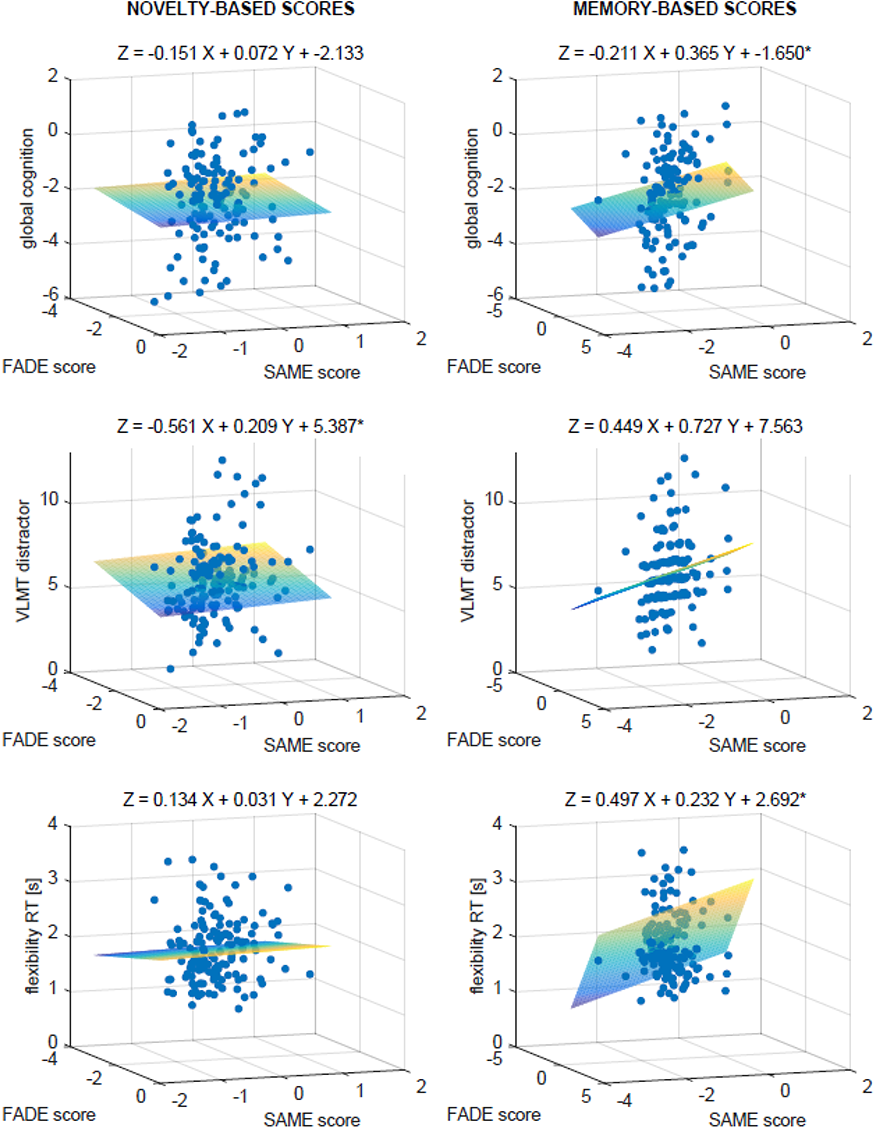
3D-Scatter plots for the regression analyses of FADE vs. SAME scores from the novelty (left) and memory contrast (right) for different neuropsychological tests as dependent variables: global cognition, the distractor list recall of the VLMT, and the reaction time in the flexibility task. * Model is significant at the .05 level. ** Model is significant at the .01 level.

**Table 3.**
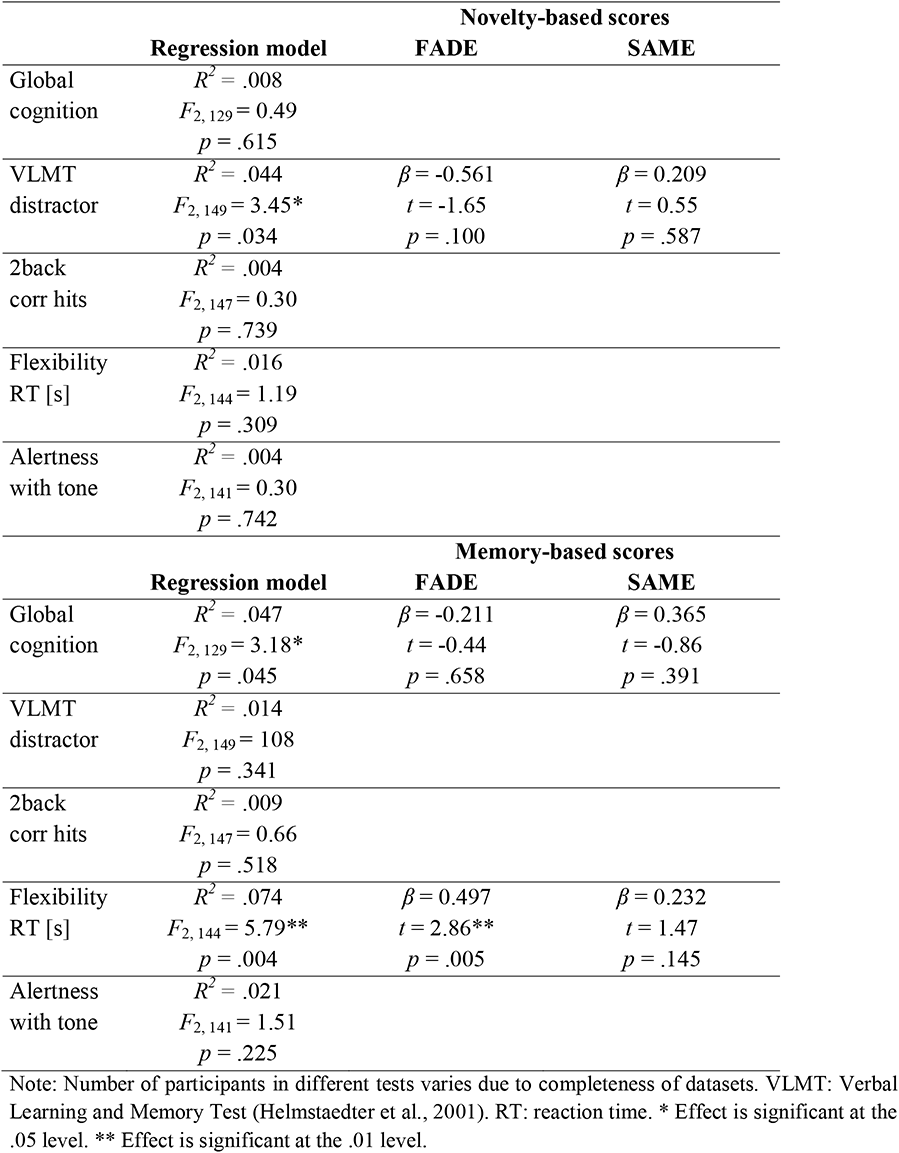
Regression analyses of the imaging scores with global cognition in older adults.

Regarding the metrics computed from the memory contrast (Table 3, lower part, and Figure 4, right side), the scores contributed meaningful information in the explanation of the RT in the flexibility task (*F*_2, 147_ = 0.66, *p* = .004), with unique impact of the FADE score (FADE: *t* = 2.86, *p* = .005, SAME: *p* = .145), and of the global cognition measure (*F*_2, 129_ = 3.18, *p* = .045), with no unique impact of either one score (FADE: *p* = .658, SAME: *p* = .391).

In the group of young adults, no significant regression results were observed (novelty scores: all *p* > .197; memory scores: all *p* > .472).

### 3.5. Correlations of the imaging scores with brain morphology

Next, we investigated the relationship of the imaging scores with age-related variability in brain morphology. In line with previous studies (Arvanitakis et al., 2016), older compared to young participants had significantly lower GMV (*t* = 6.89; *p* < .001) and higher WM lesion volumes (Mann-Whitney *U*-test: *U* = 2001.00, *p* < .001).

We observed no significant correlations between the imaging scores and WM lesion volume (Spearman’s *ρ*: all *p* > .223). For their relationship with local GMV using VBM, we detected significant correlations of the memory scores with MTL structures like the hippocampus in older adults (see Figure 5 and Table 4). The SAME memory score additionally showed correlations with local GMV in superior and inferior frontal gyrus, while the FADE memory score was additionally correlated with middle occipital gyrus GMV. *Post-hoc* analysis for the SAME memory score components revealed that the correlations were driven by the activation component while no correlations were observed for the deactivation component (see Supplementary Table S8). Furthermore, no correlations were observed for the novelty scores. The respective results from young participants can be found in Supplementary Table S9.

**Figure 5.**
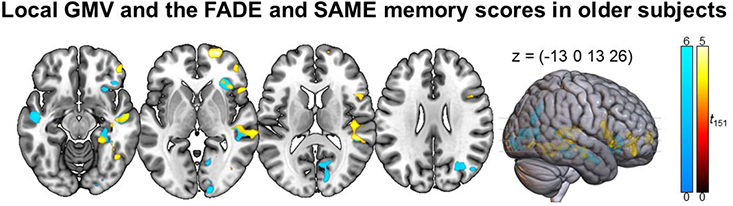
Imaging scores computed from the memory contrast and GMV using VBM. Warm colors indicate positive effects of the SAME memory score and cool colors indicate negative effects of the FADE memory score. *p* < .05, family-wise error-corrected at cluster level, cluster-defining threshold *p* < .001, uncorrected. All activation maps are superimposed on the MNI template brain provided by MRIcroGL (https://www.nitrc.org/projects/mricrogl/).

**Table 4.** Imaging scores conducted from the memory contrast and local GM volume in older participants.

|  | Hemisphere | Cluster size | Peak *t* | *p* | x, y, z |
| --- | --- | --- | --- | --- | --- |
| **FADE memory: negative effect** |  |  |  |  |  |
| Insula | R | 3162 | 5.76 | .002 | 40, 23, -4 |
|  |  |  | 4.99 |  | 31, 32, -2 |
|  |  |  | 4.71 |  | 29, 19, -7 |
| Middle temporal gyrus | L | 1627 | 4.78 | .044 | -49, -12, -16 |
|  |  |  | 3.82 |  | -51, -3, -21 |
|  |  |  | 3.22 |  | -61, -14, -10 |
| Middle temporal gyrus | R | 5674 | 4.71 | <.001 | 59, -42, 9 |
|  |  |  | 4.37 |  | 54, -10, -19 |
|  |  |  | 4.37 |  | 46, -34, 10 |
| Calcarine fissure and surrounding cortex | R | 3757 | 4.63 | .001 | 15, -68, 5 |
|  |  |  | 4.56 |  | 15, -98, -2 |
|  |  |  | 4.48 |  | 13, -85, 13 |
| Middle occipital gyrus | R | 1945 | 4.30 | .022 | 36, -69, 28 |
|  |  |  | 4.25 |  | 27, -67, 29 |
|  |  |  | 3.77 |  | 50, -71, 22 |
| Parahippocampal gyrus | R | 1736 | 4.14 | .035 | 33, -38, -8 |
|  |  |  | 3.87 |  | 27, -32, -16 |
|  |  |  | 3.74 |  | 41, -43, -18 |
| **SAME memory: positive effect** |  |  |  |  |  |
| Superior frontal gyrus, dorsolateral part | R | 1598 | 4.82 | .047 | 23, 63, 1 |
|  |  |  | 3.60 |  | 21, 62, 10 |
|  |  |  | 3.20 |  | 13, 65, 12 |
| Superior temporal gyrus, Hippocampus | R | 5503 | 4.72 | <.001 | 46, -34, 11 |
|  |  |  | 4.70 |  | 60, -41, 9 |
|  |  |  | 4.35 |  | 42, -17, -12 |
| Inferior frontal gyrus | R | 4505 | 4.68 | <.001 | 47, 19, -5 |
|  |  |  | 4.51 |  | 52, 15, 8 |
|  |  |  | 3.88 |  | 45, 42, 4 |
| Fusiform gyrus, Lingual gyrus | R | 2694 | 4.33 | .005 | 38, -47, -10 |
|  |  |  | 4.18 |  | 24, -55, -7 |
|  |  |  | 4.07 |  | 39, -72, -3 |

$p < .05$ , family-wise error-corrected at cluster level, cluster-defining threshold $p < .001$ , uncorrected.

## 4. Discussion

In previous studies, comprehensive scores reflecting memory-related fMRI activations and deactivations have been constructed as potential biomarkers for neurocognitive aging (FADE and SAME scores; E. Duzel et al., 2011; Soch et al., 2021a). Here, we aimed to further evaluate the biological relevance of these scores by investigating their relationship with performance in an extensive neuropsychological testing battery as well as brain morphological measures.

### 4.1. Neurocognitive correlates of the FADE and SAME imaging scores

While we had initially expected that, by considering both deactivation and activation deviations, the SAME score would constitute a more comprehensive or accurate measure, we found relatively few differences between the SAME and FADE scores computed from the same fMRI contrasts (i.e., novelty processing vs. subsequent memory). Instead, the fMRI contrasts had considerable influence on the relationship between the scores and indices of neurocognitive functioning. This already became evident from the inter-correlations of the imaging scores. We observed high correlations between the FADE and SAME scores derived from the same fMRI contrasts, while neither the FADE nor SAME scores computed from different fMRI contrasts correlated with each other. The implications are two-fold:

I. The FADE and SAME scores assess age-related deviation from (or similarity with) prototypical task-related activation patterns of younger participants to a comparable degree.
II. It is important to consider the functional contrast from which the scores are derived, as they appear to capture at least partly complementary information on age-related differences in cognitive function. The different contrasts reflect separable cognitive processes (novelty detection versus encoding success), and they likely capture dissociable aspects of cognitive aging, as discussed below.

Imaging scores obtained from the novelty contrast could be relatively specifically associated with performance in episodic memory tasks with unique impact of the SAME score on the explanation of memory performance for the pictures shown during fMRI scanning, of the FADE score on the WMS delayed recalls, and shared explaining variance of both scores on the recall of the VLMT distractor list. On the other hand, the imaging scores obtained from the memory contrast were significantly related to a broader set of cognitive functions, with unique impact of the SAME score on the explanation of A’ and VLMT delayed recall rates, and of the FADE score on the RTs in the flexibility task. Moreover, there was shared explaining variance of both scores when analyzing the WMS delayed recall rates and the global cognition score, which included measures of episodic memory, working memory, alertness, reaction speed, and cognitive flexibility.

One interpretation for the associations of the memory scores with cognitive (behavioral) performance beyond episodic memory could be a higher sensitivity of the memory scores towards age-related differences, as evident in the absence of an age-group effect for the FADE score computed from the novelty contrast, while the scores computed from the memory contrast showed a robust age-group differentiation (for a discussion, see Soch et al., 2021a). While the subsequent memory effect is based on the participants’ 5-point recognition-confidence ratings, the novelty contrast compares the neural responses to *de facto* novel versus highly familiarized images, not accounting for encoding success and graded confidence. Especially confidence measures are highly sensitive to aging effects (Wong et al., 2012). In our parametric design, variance attributable to both encoding success and recognition confidence was captured by the parametric subsequent memory regressor (Soch et al., 2021b). Despite the overlap of brain networks involved in novelty detection and successful episodic encoding, there are differences in detail (Maass et al., 2014, and, importantly, the memory-related brain regions contributing to the scores such as the dorsolateral and ventrolateral prefrontal cortex, the parahippocampal gyrus and MTL are not only relevant for episodic encoding but also for cognitive processes like alertness (Liu et al., 2019) or working memory (Sambataro et al., 2010; Steffener et al., 2020; Steiger et al., 2019). The novelty-related scores were significantly associated with episodic memory. Compatible with this finding, attenuated hippocampal novelty responses (E. Duzel et al., 2022) and reduced DMN deactivations during novelty processing (Billette et al., 2022) have been linked to lower memory performance in individuals at risk for AD.

### 4.2. Age-related variation in functional and structural neuroanatomy

Considering the rather specific link of the novelty-related scores with episodic memory performance in older adults, it may seem surprising that we did not observe a correlation of FADE or SAME scores with hippocampal GMV. One explanation for this could be that hippocampal volumes may correlate only moderately, if at all, with memory performance and fMRI indices of hippocampal functional integrity (E. Duzel et al., 2018; Woodard et al., 2010).

On the other hand, the FADE and SAME scores derived from the memory contrast did correlate with brain-morphometric individual differences reflecting age-related GMV loss. More specifically, we observed correlations between the memory scores and local GMV for hippocampus, parahippocampal gyrus, middle temporal gyrus and prefrontal cortex using VBM. Importantly, all of these correlations were observed in the older age group only, suggesting that they reflect individual differences related to aging rather than development or general cognitive ability. Concurrent brain-structural alterations and lower cognitive performance in aging constitute a well-replicated finding. Hedden et al. (2016) examined the relationship between age-related cognitive impairment and various brain markers (MRI and PET) and observed associations of striatal volume and WM integrity with processing speed and executive functions, and of hippocampal volume and amyloid load (as assessed with PET) with episodic memory. Considering the memory-related scores and their association with cognitive function beyond episodic memory and with brain morphology, our results are compatible with previous findings in other cohorts. Arvanitakis et al. (2016) found lower whole-brain GMV to be associated with episodic memory performance and perceptual speed. Similarly, Tsapanou et al. (2019) observed that age-related differences in episodic memory, processing speed and executive functions were associated with cortical thickness, WM hyperintensities and striatal volume. In a large cohort of over 3000 healthy participants, Zonneveld et al. (2019) reported an association of global cognition with GMV in the left amygdala, hippocampus, parietal lobule, superior temporal gyrus, insula and posterior temporal lobe. One potential advantage of our fMRI-based scores becomes evident from the recent observation that the scores may be superior to structural MRI data – and also resting-state fMRI – in the prediction of memory performance in older adults (Soch et al., in press). Future investigations should therefore explore the possibility that fMRI-based markers may be suitable as a predictor of cognitive functioning, even when age-related structural changes are not (yet) observable.

### 4.3. Deactivation of the Default Mode Network and cognitive function in old age

While the influence of the underlying contrast (novelty vs. memory) generally outweighed the effects of score type (FADE vs. SAME), in the few cases where the SAME compared to the FADE score did show unique associations with additional functions (e.g., A’, VLMT delayed recall performance as well as local GMV in frontal cortex), these associations were mainly driven by the deactivation component of the SAME score.

This pattern can likely be attributed to the construction of the SAME score, also including age-dependent differences in functional deactivation patterns, while the FADE score only relies on activation differences. Brain regions that showed prominent deactivations during successful memory encoding in the young participants included a network centered around the brain’s midline that has previously been referred to as the DMN (Raichle, 2015). This observation is in line with a frequently cited meta-analysis by Maillet and Rajah (2014), who found age-related differences in encoding-related processes encompassing under-recruitment of occipital, parahippocampal, and fusiform cortex, but over-recruitment of DMN regions including the medial prefrontal cortex (mPFC), precuneus, and left inferior parietal lobe in older adults. In the current study, the correlation of the SAME memory score with global cognition could be primarily accounted for by the deactivation component, which may, at least in part, reflect an older individual’s general ability to suppress ongoing DMN activation during attention-demanding tasks. In line with this interpretation, reduced DMN deactivation has also been associated with lower working memory performance in older adults (Sambataro et al., 2010), and a meta-analysis revealed that reduced DMN deactivation in old age can be observed across a variety of cognitive tasks (H. J. Li et al., 2015). On the other hand, several authors discuss the role of the DMN as a potential cognitive resource in older adults (Billette et al., 2022; Colangeli et al., 2016), which should be further addressed in future studies (see Supplementary Discussion).

### 4.4. A potential role for the mesolimbic dopamine system in successful aging

Among the scores investigated here, the SAME score from the novelty contrast stood out by showing a positive correlation with voxel-wise activations not only for the novelty contrast (Figure S1), but also for the subsequent memory contrast (Figure 2). Notably, the peak of this correlation was found in the striatum, a core output region of the midbrain dopaminergic nuclei. Previous studies have implicated the dopaminergic midbrain in successful encoding in young adults (Adcock et al., 2006; Schott et al., 2006; Wittmann et al., 2005). In older adults, striatal dopamine D2 receptor binding has been related to hippocampal-striatal functional connectivity and memory performance (Nyberg et al., 2016). Importantly, novelty can induce midbrain activations (Bunzeck & Duzel, 2006; Schott et al., 2004), and structural integrity of the midbrain has been related to both midbrain and hippocampal novelty responses (Bunzeck et al., 2007) and to memory performance in older adults (S. Duzel et al., 2008). Düzel et al. (2010) proposed the NOMAD model which suggests that novelty-related increase of mesolimbic dopaminergic activity promotes exploratory behaviour and ultimately memory performance in older adults. In line with this framework, our results suggest that preserved patterns of brain responses to novelty may be related to increased activity of mesolimbic dopaminergic structures during successful memory formation in aging.

### 4.5. Implications for clinical research

Quantification of neurocognitive aging and early identification of individuals at risk for accelerated cognitive decline may help to ultimately develop targeted early interventions to improve cognitive functioning in older adults. Especially early lifestyle interventions, tackling physical exercise, nutrition, and to some degree cognitively demanding tasks, can be helpful to preserve healthy aging (Bishop et al., 2010; Franke & Gaser, 2019; Stern, 2012; Whitty et al., 2020). However, an accurate assessment of cognitive, but also neurophysiological decline poses a major challenge due to the complexity of brain processes and functions, as well as the non-linear acceleration of cognitive decline (Vinke et al., 2018).

As the present study was directed at the association between fMRI-based markers for network dysfunction and neurocognitive functioning in healthy older adults, the next step should be to test our scores in (pre-)clinical populations where dysfunctions of successful-encoding and novelty networks are prominent and may even precede neuropsychological impairment or brain morphometric changes like atrophy (Zhou & Seeley, 2014). With respect to AD, the scores may be of interest in the investigations of individuals with mild cognitive impairment (MCI), a clinical condition with considerable diagnostic and prognostic uncertainty, such that higher accuracy in diagnosis would be of high clinical value. In older adults with MCI and related risk states like subjective cognitive decline (SCD; Jessen et al., 2022), various biomarkers have been assessed for their potential clinical applicability. However, thus far, task-based fMRI has largely focused on dysfunctional hippocampal activity (Marquez & Yassa, 2019). The mediocre test-retest reliability of voxel-wise task-based fMRI has called into question its utility as a biomarker (Elliott et al., 2020). The reductionist single-value scores of age-related whole-brain fMRI activation (and deactivation) patterns described here may prove more reliable. In this context, it is of importance that in recent studies with older participants at risk for AD, researchers have often employed novelty rather than subsequent memory contrasts, owing to the lack of successfully encoded items in individuals with pronounced memory impairment (Billette et al., 2022; E. Duzel et al., 2018; E. Duzel et al., 2022). Our observation that the novelty-related scores, particularly the FADE novelty score, show relatively strong and specific correlations with tests of hippocampus-dependent memory, support the validity of this approach. It may nevertheless be of interest what the memory-related scores, and particularly the SAME memory score, signify in memory-impaired individuals. They may, for example, prove a useful tool in the assessment of cognitive impairment beyond the memory domain or in atypical presentations of pre-clinical dementia. The scores may also help to better understand and define "healthy aging" on a theoretical level and could facilitate the laborious screening of high-risk patients for pharmacological studies or may be combined with tau-or amyloid-PET (Billette et al., 2022) as a potential biomarker assessment at the clinical level.

### 4.6. Limitations

We analyzed data from a cross-sectional cohort of healthy older adults. As the measured variables deteriorate with age, future longitudinal studies would be needed to better understand the relationship between functional and structural imaging as well as neuropsychological performance changes as ageing progresses and eliminate age-related confounds in cross-sectional studies (Elliott, 2020; Xing, 2021).

Another limitation is, that the maximum explained variance was an R-squared of .114 for the explanation of the WMS delayed recalls, suggesting that around 90 % of the variation in cognitive functions are not explained by the single-value scores.

Importantly the observed associations were only apparent in the group of older adults and not in the group of young participants. Further analysis capturing the whole lifespan and especially an appropriate sized group of middle-aged participants are needed to further evaluate the predictive values of the singe-value fMRI scores.

### 4.7. Conclusion

Our results provide novel brain-behavior associations of single-value fMRI-based scores with cognitive ability in older adults. They further suggest that the scores provide complementary information with respect to relatively selective impairment of hippocampal function versus general cognitive ability and local GMV loss in old age. Future research should address their utility and predictive value in (pre-)clinical populations like AD (risk states).

## Supporting information

Supplementary Methods, Results, Figures, and Tables

## 6. Statements

## 6.1. Acknowledgments

We are grateful to Herta Flor and Shari Wiseman for valuable comments on the manuscript. We thank Adriana Barman, Marieke Klein, Kerstin Möhring, Katja Neumann, Ilona Wiedenhöft, and Claus Tempelmann for assistance with MRI data acquisition.

## 6.2. Data Availability Statement

Due to data protection regulations, sharing of the entire data set underlying this study in a public repository is not possible. We have previously provided GLM contrast images as a NeuroVault collection (https://neurovault.org/collections/QBHNSRVW/) and MATLAB code for imaging scores as a GitHub repository (https://github.com/JoramSoch/FADE_SAME) for an earlier article using the same dataset Soch et al., 2021a. Access to de-identified raw data will be provided by the authors upon reasonable request.

