## Supplementary Methods, Results, Figures, and Tables for "Single-value scores of memory-related brain activity reflect dissociable neuropsychological and anatomical signatures of neurocognitive aging"

### 1. Supplementary Methods

#### 1.1. Participants

**Table S1. Demographics of young and older subjects.**

| | young subjects | older subjects | statistics |
|---|---|---|---|
| N | 106 | 153 | – |
| mean age ± SD, age range | 24.12 ± 4.00 yrs, 18-35 yrs | 64.04 ± 6.74 yrs, 51-80 yrs | $t$ = -59.66, $p$ < .001 |
| gender ratio (m/f) | 47/59 | 59/94 | $\chi^2$ = 0.87, $p$ = .352 |
| MMSE mean performance ± SD | – | 28.93 ± 0.99 (range: 26-30) | – |
| educational status (with/without "Abitur") | 100/6 | 72/81 | $\chi^2$ = 62.75, $p$ < .001 |
| MWT-B mean hits ± SD | 26.81 ± 3.14 (range: 21-35) | 30.33 ± 3.25 (range: 19-36; 1 missing) | $z$ = -8.11, $p$ < .001 |
| ethnic composition (European/other) | 104/2 | 153/0 | $\chi^2$ = 2.91, $p$ = .088 |
| ApoE genotype | | | $\chi^2$ = 3.03, $p$ = .696 |
| E2/E2 | 1 | 3 | |
| E2/E3 | 15 | 26 | |
| E2/E4 | 4 | 4 | |
| E3/E3 | 60 | 94 | |
| E3/E4 | 24 | 24 | |
| E4/E4 | 2 | 2 | |
| medication | | | – |
| oral antidiabetics | 0 | 1 | |
| antihypertensives | 2 | 49 | |
| statins | 0 | 4 | |
| thyroid hormone | 1 | 18 | |
| contraceptives and/or hormone replacement | 28 | 7 | |
| endocrine-related surgery: history of | | | – |
| thyroidectomy | 0 | 2 | |
| oophorectomy | 0 | 2/3 uni-/bilateral | |

*Demographic information for the two age groups, along with statistics from a two-sample t-test (mean age), chi-squared tests (gender ratio, educational status, ethnic composition, and ApoE genotype) and a Mann-Whitney U-test (MWT-B hits). Abbreviations: N = sample size, SD = standard deviation, yrs = years, m = male, f = female, MMSE = Mini-Mental State Examination (Creavin et al., 2016; Folstein et al., 1975), MWT-B = multiple choice vocabulary intelligence test ("Mehrfachwahl-Wortschatz-Intelligenztest", Lehrl, 2005). "Abitur" is the German equivalent of a high school graduation certificate (university entrance level). Age, gender, ethnicity, education, medication, and history of endocrine-related surgery according to self-report in a health questionnaire. For details on ApoE genotyping, see Soch et al. (2021).*

**1.2. Neuropsychological testsVerbal Learning and Memory Test (VLMT)**

An adapted version of the Verbal Learning and Memory Test (VLMT; Helmstaedter et al., 2001) was used to test different aspects of verbal memory functions. In the first part, a 15-noun word list (list A) was shown to the participants (presentation duration 1 s, fixation 2 s). This procedure was repeated five times and after each repetition trial, subjects were requested to freely recall as many words as possible and write them down. To assess the learning ability, the number of correctly reproduced items was added up across the five trials. Subsequently, a second list of 15 different nouns (list B) was presented, again followed by a free recall test. This second list served as an interference for a delayed memory test of list A. Immediately after recall of list B, the participants were again asked to recall list A. Delayed recall of list A was measured 30 min after its immediate recall. The original VLMT moreover included a recognition list that was not used in the current study. Instead, memory for the words from list A was again assessed after a latency period of one day (Assmann et al., 2021; Barman et al., 2014). We further assessed different aspects of memory performance: the number of correctly named items of repetition trials 1 to 5 (sum score), of distractor list B, of list A directly after the distractor list, and at the delayed recalls after 30 minutes and one day. This served the purpose of capturing memory performance as a function of the different retention intervals.

**1.2.2. Logical Memory subtest from the Wechsler Memory Scale (WMS)**

To assess logical memory, we used a subtest from the German version of Wechsler Memory Scale test battery (WMS; Härting et al., 2000). In this task, participants listened to two different short stories via headphones. They were asked to memorize these as accurately as possible and write them down immediately after. After a 30-minute delay as well as a latency period of one day, free recall of both stories was assessed. Both stories were divided into a total of 25 short segments. Evaluation of memory performance was based on precise evaluation guidelines by two independent raters. Each segment was scored with one point if retrieved correctly and the points for the two stories were summed up. Mean values of the two raters were used for further analysis.

##### 1.2.3. Alertness subtest from the Test Battery for Attention (TAP)

The alertness task is a subtest from the Test Battery for Attention (TAP; Zimmermann & Fimm, 1993). It is divided into tonic (stable and general alertness) and phasic alertness (short-term and rapid increase in attention). In the task a cross appeared on the screen at randomly varying intervals (presentation duration 0.5 s, fixation 3.5-4s) and participants were instructed to respond to the cross as quickly as possible by pressing a button. In the phasic alertness trials, a warning tone as a cue preceded the cross. The subjects performed two blocks per condition with 20 trials each block. The reaction time (RT) in tonic and phasic alertness trials was used for analysis.

##### 1.2.4. Flexibility subtest from the Test Battery for Attention (TAP)

Executive functions were examined using, among others, a flexibility paradigm, which is another subtest from the TAP (Zimmermann & Fimm, 1993). It assesses the ability to change the focus of attention. In the TAP’s flexibility paradigm, competing stimuli are presented simultaneously on the screen right and left of the fixation point. We used the complex verbal variant, where the competing stimuli consisted of a letter and a number and the target category alternates. From one presentation to the other, the target changes either from letter to number and vice versa. Participants were instructed to press the key on the side of the displayed target stimulus as fast as possible. Dependent variables were RT for correct responses and the percentage of errors.

##### 1.2.5. Flanker task

We used a modified version of the Flanker task that is one of the classic paradigms for studying interference processing and cognitive control (Eriksen & Eriksen, 1974). The subject's task was to determine the direction of a center arrow in a horizontally line of 5 arrows by pressing a key (presentation duration 0.5 s). In the congruent condition, flanking arrows pointed in the same direction as the target arrow and in the incongruent condition, the flanking arrows pointed in the opposite direction. Additionally, baseline trials (lines as distractors) and no-go trials (appearance of a circle as stop signal) were included. Ten runs with 30 trials each were conducted. The analyzed characteristic of the Flanker task was the so-called congruency effect, i.e. the difference in average RT between correct incongruent and correct congruent go trials.

###### 1.2.6. N-Back task

The n-back paradigm was used to test working memory functions (Gevins & Cutillo, 1993). Participants were presented with a sequence of stimuli and instructed to respond whenever the current stimulus corresponded to the stimulus presented *n* steps earlier. The factor *n* represents the degree of difficulty (Kirchner, 1958). The task was performed including 1-back, 2-back, and 3-back conditions (trial presentation duration 1 s, inter-trial interval 0.25 s). In the 1-back condition, the subject was to respond, if the item presented was identical to the item presented immediately before. Successful completion of the 3-back condition required a response when the stimulus presented matched the stimulus that came three trials before the current trial. In the version used here, single letters were presented in the center of the computer screen. Each n-back condition occurred four times (12 blocks in total). In each block, 24 letters were presented with four correct responses. The n-back task was evaluated in terms of corrected hit rates (hit rate minus false alarm rate) and in terms of RTs for hits in 1-back, 2-back, and 3-back trials.

#### 1.3. Outlier detection

**Table S2. Variables and borders for removal of extreme outliers in older and young participants**

| test | variables | extreme outlier removal for older participants | extreme outlier removal for young participants |
|---|---|---|---|
| Verbal Learning and Memory Test (VLMT) | number of correctly named words of | | |
| | - repetitions of list A (sum score) | - | - |
| | - distractor list B | - | - |
| | - recall of list A | - | - |
| | - 30-min delayed recall of list A | - | - |
| | - one-day delayed recall of list A | - | - |
| Logical Memory subtest from the WMS | number of story details retrieved at: | | |
| | - immediate recall | - | - |
| | - 30-min delayed recall | - | - |
| | - one-day delayed recall | - | - |
| Alertness subtest from the TAP | reaction on the appearance of a cross: | | |
| | - RT in trials with cue tone | - | - |
| | - accuracy in trials with cue tone | < 80% | < 80% |
| | - RT in trials without cue tone | > 612ms | - |
| | - accuracy in trials without cue tone | < 97% | < 97% |
| Flexibility subtest from the TAP | switching attention between targets: | | |
| | - error rate | - | > 24% |
| | - RT | - | - |
| Flanker task | incongruent vs. congruent trials: | | |
| | - accuracy in congruent trials | < 85 % | < 90 % |
| | - RT in congruent trials | > 1140 ms | > 814 ms |
| | - accuracy in incongruent trials | - | < 45 % |
| | - RT in incongruent trials | > 1766 ms | - |
| N-Back task | responses on reoccurring letters: | | |
| | - 1-back corrected hit rate | < 12 % | < 76 % |
| | - 1-back RT | > 871 ms | > 696 ms |
| | - 2-back corrected hit rate | - | < -64 |
| | - 2-back RT | < 208 ms | - |
| | - 3-back corrected hit rate | - | < -132 % |
| | - 3-back RT | > 1397 ms | - |

*Classification as extreme outliers based on the interquartile range (IQR; x > 3rd quartile + 3*IQR, x < 1st quartile – 3*IQR). RT: reaction time. WMS: Wechsler Memory Scale (Härting et al., 2000). TAP: Test Battery for Attention (Zimmermann & Fimm, 1993). VLMT: Verbal Learning and Memory Test (Helmstaedter et al., 2001).*

### 2. Supplementary Results

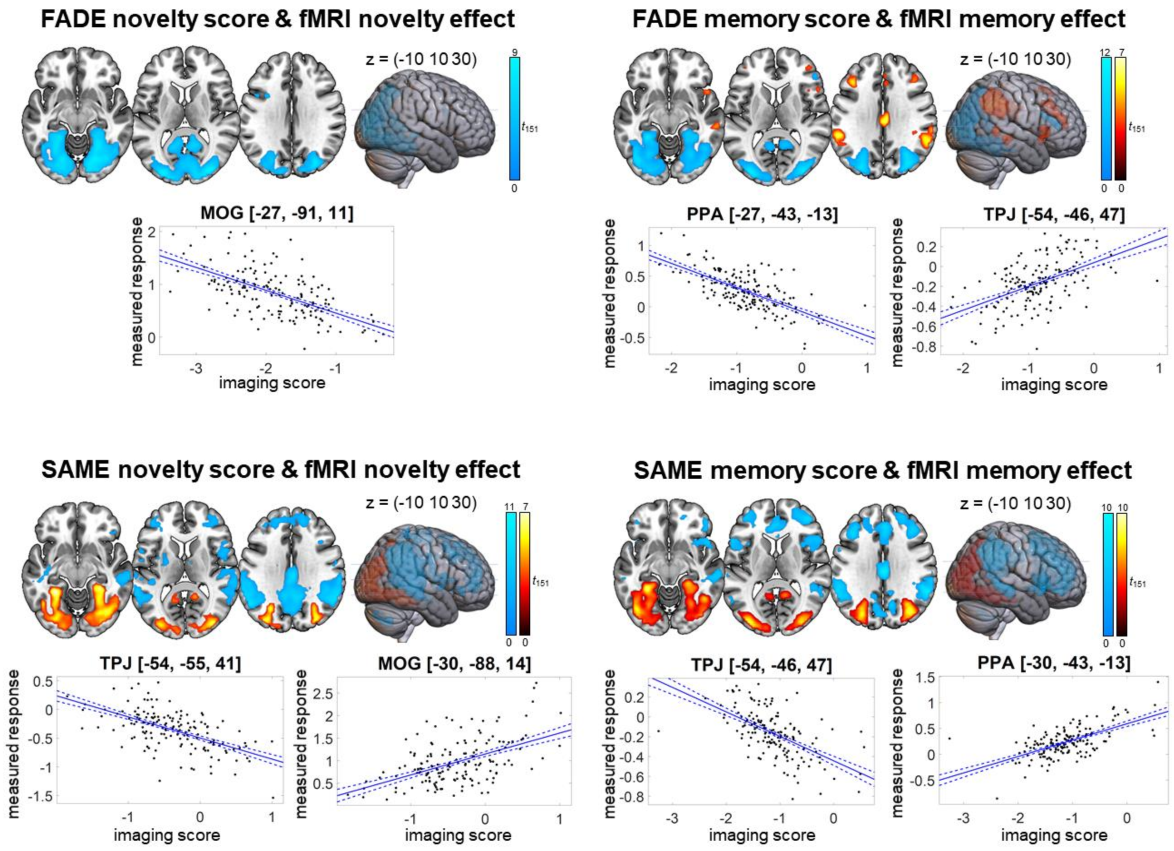


***Figure S1.*** *Imaging scores and fMRI effects (novelty effect and subsequent memory effect) in older participants. Warm colors indicate positive effects and cool colors indicate negative effects. p < .05, family-wise error-corrected at cluster level, cluster-defining threshold p < .001, uncorrected. MOG: middle occipital gyrus, PPA: parahippocampal place area, TPJ: temporo-parietal junction. All activation maps are superimposed on the MNI template brain provided by MRIcroGL (https://www.nitrc.org/projects/mricrogl/).*

**Table S3. Imaging scores and fMRI effects in older participants: FADE novelty score & fMRI novelty effect**

| | Hemisphere | Cluster size | peak *t* | *p* | x y z |
|---|---|---|---|---|---|
| **negative effect** | | | | | |
| Lingual gyrus | R | 4562 | 9.61 | <.001 | 21 -73 -7 |
| | | | 9.14 | | -27 -91 11 |
| | | | 8.80 | | 30 -79 26 |
| Inferior frontal gyrus | L | 33 | 4.74 | .048 | -30 8 32 |
| | | | 4.36 | | -45 11 26 |
| Inferior parietal gyrus | L | 58 | 4.23 | .003 | -27 -52 53 |
| | | | 4.22 | | -21 -58 59 |
| | | | 3.53 | | -15 -58 50 |

*$p < .05$, family-wise error-corrected at cluster level, cluster-defining threshold $p < .001$, uncorrected. MOG: Middle occipital gyrus.*

**Table S4. Imaging scores and fMRI effects in older participants: SAME novelty score & fMRI novelty effect**

| | Hemisphere | Cluster size | peak *t* | *p* | x y z |
|---|---|---|---|---|---|
| **negative effect** | | | | | |
| Angular gyrus | R | 1555 | 11.28 | <.001 | 54 -58 38 |
| | | | 9.45 | | 48 -58 47 |
| | | | 8.60 | | 60 -43 35 |
| Angular gyrus | L | 2901 | 9.83 | <.001 | -51 -61 35 |
| | | | 9.58 | | 9 -52 35 |
| | | | 8.76 | | -54 -55 41 |
| Superior frontal gyrus | L | 1310 | 6.61 | <.001 | -12 26 56 |
| | | | 6.48 | | -36 26 38 |
| | | | 6.34 | | 15 47 41 |
| Suppl. motor area | R | 69 | 5.60 | .001 | 6 14 65 |
| | L | | 4.29 | | -9 14 65 |
| | L | | 3.84 | | -21 5 68 |
| Cerebellum | L | 67 | 5.32 | .001 | -30 -79 -37 |
| | | | 5.22 | | -24 -85 -37 |
| | | | 3.85 | | -39 -73 -40 |
| Cerebellum | R | 49 | 5.27 | .008 | 42 -64 -43 |
| | | | 4.28 | | 36 -76 -40 |
| | | | 3.97 | | 33 -82 -34 |
| Postcentral gyrus | R | 142 | 5.01 | <.001 | 15 -40 74 |
| | | | 4.42 | | 33 -37 65 |
| | | | 4.40 | | 33 -28 65 |
| Thalamus | R | 42 | 4.75 | .017 | 12 -7 17 |
| | | | 3.93 | | 0 -1 5 |
| Inferior frontal gyrus | L | 62 | 4.51 | .002 | -54 5 8 |
| | | | 4.02 | | -57 -1 2 |
| | | | 3.56 | | -57 14 20 |
| Caudate nucleus | L | 90 | 4.44 | <.001 | -18 -7 23 |
| | | | 4.33 | | -24 -7 11 |
| | | | 3.70 | | -12 -13 14 |
| Superior frontal gyrus | R | 34 | 4.33 | .041 | 3 53 2 |
| **positive effect** | | | | | |
| MOG | R | 2358 | 7.31 | <.001 | 33 -82 20 |
| | | | 7.24 | | 36 -85 5 |
| | | | 7.20 | | -30 -88 14 |
| Precuneus and PCC | R | 55 | 5.47 | .004 | 15 -55 14 |
| | | | 4.66 | | 12 -49 8 |
| Calcarine fissure and | L | 40 | 4.77 | .021 | -6 -52 5 |
| surrounding cortex | | | 4.03 | | -15 -55 11 |

*$p < .05$, family-wise error-corrected at cluster level, cluster-defining threshold $p < .001$, uncorrected. MOG: Middle occipital gyrus, TPJ: Temporoparietal junction.*

**Table S5. Imaging scores and fMRI effects in older participants: FADE memory score & fMRI memory effect**

| | Hemisphere | Cluster size | peak *t* | *p* | x y z |
|---|---|---|---|---|---|
| **negative effect** | | | | | |
| MOG | L | 1896 | 11.89 | <.001 | -33 -88 14 |
| | | | 10.83 | | -30 -85 23 |
| | | | 10.31 | | -27 -43 -13 |
| MOG | R | 1986 | 10.64 | <.001 | 36 -85 17 |
| | | | 10.34 | | 33 -79 23 |
| | | | 10.28 | | 36 -88 8 |
| Inferior frontal gyrus | R | 66 | 4.99 | .002 | 51 32 17 |
| | | | 4.72 | | 51 38 8 |
| | | | 4.66 | | 45 20 23 |
| **positive effect** | | | | | |
| TPJ | L | 336 | 7.33 | <.001 | -54 -46 47 |
| | | | 6.79 | | -57 -40 29 |
| | | | 4.78 | | -42 -49 56 |
| Supramarginal gyrus | R | 420 | 7.20 | <.001 | 57 -40 44 |
| | | | 6.93 | | 57 -52 38 |
| | | | 4.81 | | 45 -49 53 |
| Posterior cingulate gyrus | R | 136 | 6.54 | <.001 | 3 -22 32 |
| | | | 3.64 | | 0 -34 23 |
| | | | 3.37 | | -3 -7 35 |
| Middle frontal gyrus | L | 105 | 5.94 | <.001 | -39 29 32 |
| | | | 3.68 | | -42 44 14 |
| Middle temporal gyrus | R | 41 | 5.16 | .02 | 63 -28 -7 |
| Inferior frontal gyrus | R | 93 | 5.06 | <.001 | 48 20 -1 |
| | | | 4.16 | | 51 14 -7 |
| | | | 3.68 | | 39 20 8 |
| Middle frontal gyrus | R | 139 | 4.94 | <.001 | 39 38 23 |
| | | | 4.80 | | 39 23 38 |
| | | | 4.14 | | 42 47 14 |
| Middle temporal gyrus | R | 38 | 4.43 | .028 | 57 -43 2 |
| Superior frontal gyrus | R | 38 | 4.06 | .028 | 15 26 53 |
| | | | 4.01 | | 18 17 59 |
| Anterior cingulate gyrus | R | 35 | 3.86 | .039 | 3 26 29 |
| | | | 3.81 | | 3 35 32 |

*$p < .05$, family-wise error-corrected at cluster level, cluster-defining threshold $p < .001$, uncorrected. MOG: Middle occipital gyrus, PPA: Parahippocampal place area, TPJ: Temporoparietal junction.*

**Table S6. Imaging scores and fMRI effects in older participants: SAME memory score & fMRI memory effect**

| | Hemisphere | Cluster size | peak *t* | *p* | x y z |
|---|---|---|---|---|---|
| **negative effect** | | | | | |
| Inferior parietal gyrus | L | 792 | 10.18 | <.001 | -54 -55 41 |
| | | | 8.69 | | -54 -46 47 |
| | | | 7.78 | | -57 -37 44 |
| Angular gyrus | R | 1005 | 10.17 | <.001 | 57 -52 38 |
| | | | 9.39 | | 51 -58 41 |
| | | | 8.56 | | 57 -40 44 |
| Median cingulate and paracingulate gyri | R | 3466 | 10.01 | <.001 | 3 -22 32 |
| | | | 8.04 | | -39 29 32 |
| | | | 7.79 | | 39 29 35 |
| Precuneus | R | 386 | 6.85 | <.001 | 12 -61 38 |
| | | | 6.77 | | -6 -70 35 |
| | | | 5.48 | | 12 -49 32 |
| Inferior frontal gyrus | L | 235 | 5.90 | <.001 | -54 17 8 |
| | | | 5.29 | | -51 14 -1 |
| | | | 4.68 | | -39 14 5 |
| Middle temporal gyrus | L | 44 | 5.07 | .014 | -63 -37 -4 |
| Cerebellum | L | 56 | 4.70 | .004 | -39 -52 -46 |
| | | | 4.54 | | -45 -61 -37 |
| | | | 3.27 | | -39 -67 -46 |
| **positive effect** | | | | | |
| MOG | L | 1507 | 10.41 | <.001 | -33 -88 17 |
| PPA | | | 9.57 | | -30 -43 -13 |
| | | | 8.87 | | -36 -88 8 |
| MOG | R | 1462 | 8.93 | <.001 | 36 -73 26 |
| | | | 8.83 | | 36 -85 20 |
| Calcarine fissure and | | | 8.49 | | 15 -52 17 |
| surrounding cortex | L | 68 | 5.32 | .001 | -15 -49 5 |
| | | | 5.26 | | -12 -55 17 |

*$p < .05$, family-wise error-corrected at cluster level, cluster-defining threshold $p < .001$, uncorrected. MOG: Middle occipital gyrus, PPA: Parahippocampal place area, TPJ: Temporoparietal junction.*

**Table S7. Imaging scores and fMRI effects in older participants: SAME novelty score & fMRI memory effect**

| | Hemisphere | Cluster size | peak *t* | *p* | x y z |
|---|---|---|---|---|---|
| **positive effect** | | | | | |
| Striatum | R | 62 | 5.46 | .003 | 9 11 2 |
| | | | 4.25 | | 0 -1 5 |
| | | | 3.95 | | 6 2 -4 |
| MOG | L | 103 | 4.97 | <.001 | -36 -79 29 |
| | | | 4.13 | | -33 -64 29 |
| | | | 3.76 | | -33 -67 38 |
| Precuneus | L | 70 | 4.41 | .001 | -3 -61 17 |
| | | | 3.83 | | 0 -52 11 |
| | | | 3.82 | | -6 -55 26 |

*p < .05, family-wise error-corrected at cluster level, cluster-defining threshold p < .001, uncorrected. MOG: Middle occipital gyrus.*

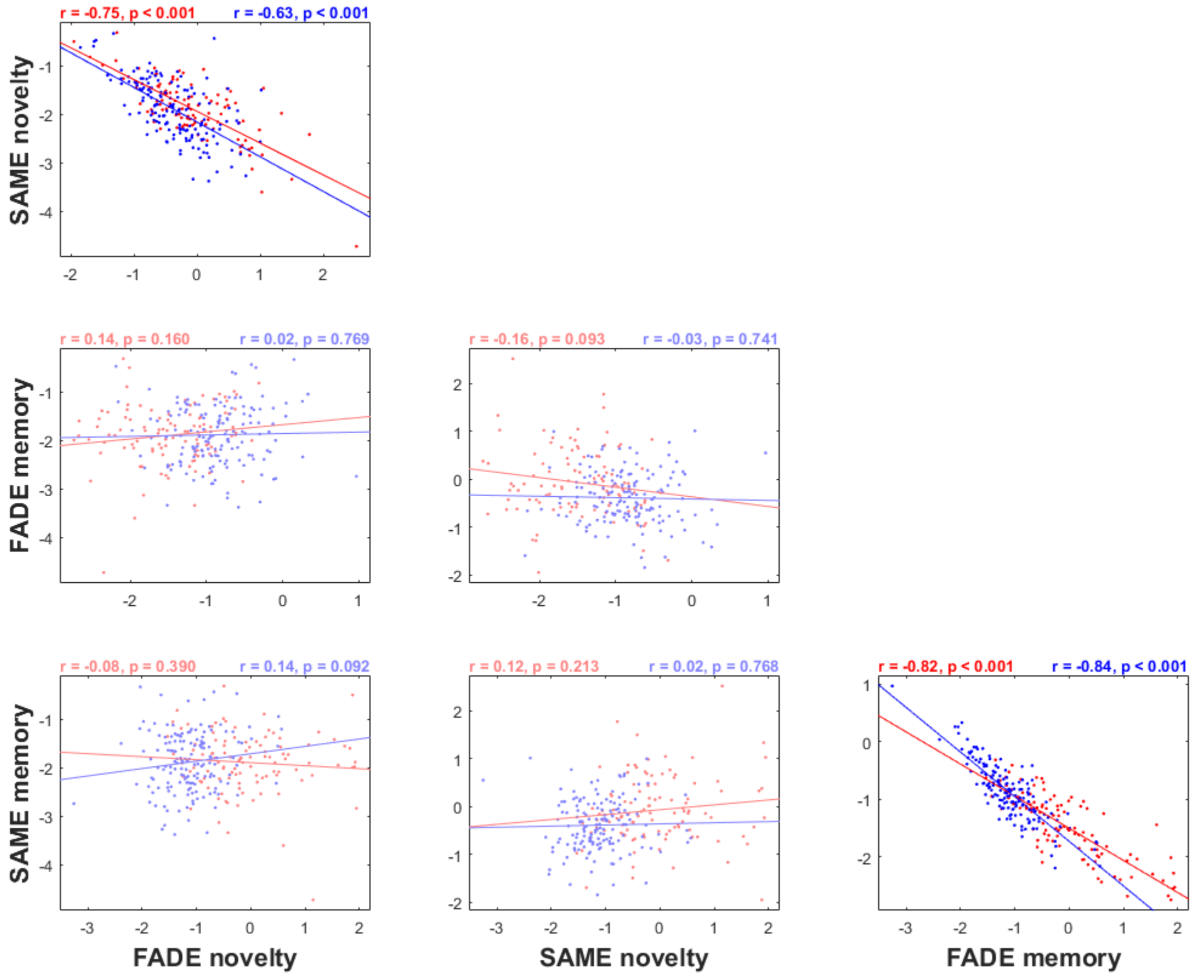


***Figure S2.*** *Pearson correlations between the FADE and SAME imaging scores conducted from the novelty and memory fMRI contrasts, separated by age group (red: young, blue: older subjects). Each dot represents one participant. Highlighted: correlation is significant at the 0.05 level (two-tailed).*

**Table S8. Activation component of the SAME score computed from the memory contrast and local GM volume in older participants**

| | Hemisphere | Cluster size | peak *t* | *p* | x y z |
|---|---|---|---|---|---|
| **SAME memory activation component: positive effect** | | | | | |
| Calcarine fissure and surrounding cortex | R | 4663 | 5.07 | .001 | 10 -67 10 |
| | | | 4.21 | | 30 -41 -6 |
| | | | 4.12 | | 12 -67 -2 |
| Inferior frontal gyrus | R | 2135 | 5.02 | .033 | 48 15 13 |
| | | | 4.75 | | 45 21 -5 |
| | | | 3.62 | | 56 24 -5 |
| Supramarginal gyrus | R | 1851 | 4.64 | .041 | 59 -13 26 |
| | | | 4.48 | | 53 -23 32 |
| | | | 3.86 | | 51 -17 37 |

*$p < .05$, family-wise error-corrected at cluster level, cluster-defining threshold $p < .001$, uncorrected.*

**Table S9. FADE score computed from the memory contrast and local GM volume in young participants**

| | Hemisphere | Cluster size | peak *t* | *p* | x y z |
|---|---|---|---|---|---|
| **FADE memory: positive effect** | | | | | |
| Insula | L | 1659 | 4.55 | .039 | -40 9 4 |
| | | | 3.89 | | -38 9 17 |
| | | | 3.91 | | 7 -78 -23 |
| Cerebellum | R | 1584 | 3.72 | .046 | 17 -80 -23 |
| | | | 3.41 | | 34 -85 -36 |

*$p < .05$, family-wise error-corrected at cluster level, cluster-defining threshold $p < .001$, uncorrected.*

#### 3. Supplementary Discussion

The interpretation that the correlation of the SAME-memory score with general cognitive ability merely reflects an unspecific reduction of DMN deactivation may be challenged by the observation that older adults, and particularly those with more pronounced age-related memory decline, even exhibit task-related activations above baseline in DMN structures during successful memory encoding (Maillet & Rajah, 2014). As suggested by Maillet and Rajah, this might reflect older adults' tendency to employ mnemonic strategies that engage self-referential processing or autobiographical memory retrieval – processes that engage the DMN also in young adults (Schilbach et al., 2012; Soch et al., 2017; Warren et al., 2018). In line with the interpretation of the DMN as a potential cognitive resource in aging, a meta-analysis by Colangeli et al. (2016) revealed that activation in bilateral medial frontoparietal areas (including precuneus, anterior cingulate and superior frontal gyrus) was associated with better performance in cognitive tasks in healthy and pathological aging, which the authors attributed to a higher cognitive reserve of the participants. However, in that meta-analysis, a cluster within the canonical DMN (pregenual/subgenual anterior cingulate) was only seen in cognitively impaired individuals. The clusters found in healthy older adults were found in a more dorsal part of the anterior cingulate, which is more likely part of the salience network than the DMN (Margulies & Uddin, 2019), and the dorsal part of the precuneus, which is involved in spatial cognition, including memory formation of complex spatial information (Schott et al., 2019). The role of the DMN as a cognitive resource may thus be most relevant in older individuals with cognitive impairment, compatible with a recently observed inverse U-shape function of memory-related precuneus activation in older adults with Alzheimer's disease risk states (Billette et al., 2022).
